# Neutrophils impair B cell differentiation via mitochondrial and lipid metabolism in lupus

**DOI:** 10.64898/2026.02.03.703615

**Authors:** Hannah F. Bradford, Guillem Montamat, Leire Goicoechea Barrenechea, Antonella Spinazzola, David M Lowe, Alan D Salama, Claudia Mauri, Marilina Antonelou

## Abstract

Systemic lupus erythematosus (SLE) is characterised by aberrant neutrophil activation and pathogenic B-cell responses that drive organ damage, particularly in lupus nephritis (LN). Here, we identify neutrophil-derived mitochondrial DNA (mtDNA) as a metabolic driver of B-cell dysregulation in LN. Using single-cell transcriptomics of human blood and kidney tissue together with functional co-culture assays, we show that neutrophils from patients with active LN release extracellular traps enriched in mtDNA that induce mitochondrial oxidative stress in B cells. NET-derived mtDNA suppresses IL-10–producing regulatory B cells while promoting pro-inflammatory and plasmablast differentiation through nucleic acid–dependent signalling. Mechanistically, oxidative stress drives maladaptive NADPH utilisation, diverting NADPH from cholesterol biosynthesis toward antioxidant defence, resulting in altered lipid trafficking, mitochondrial dysfunction, and impaired regulatory B-cell differentiation. Consistent with this model, kidney B cells from patients with LN exhibit transcriptional signatures of disrupted redox and lipid metabolism, and NADPH deficiency is associated with reduced regulatory B-cell differentiation in humans. These findings identify a neutrophil-driven metabolic checkpoint that governs B-cell fate in lupus nephritis.

## Introduction

Systemic lupus erythematosus (SLE) is a multisystem autoimmune disease characterised by abnormal neutrophil activation and autoreactive B cell responses. These responses drive the production of autoantibodies against neutrophil nuclear components, leading to circulating immune complex formation and subsequent deposition, causing small-vessel inflammation in target organs including the skin, the joints and the kidneys. Lupus nephritis (LN), a severe manifestation affecting up to 50% of SLE patients, is a leading predictor of poor prognosis with a 30% risk of progressing to end-stage renal failure requiring dialysis or transplantation^1^. The risk of kidney failure has remained largely unchanged over the past 30 years, underscoring the urgent need for more targeted therapeutic approaches.

B cell depletion with rituximab, a CD20-targeting monoclonal antibody, induces remission in only about half of patients with SLE. Nonetheless, it has highlighted the pathogenic role of B cells in the development of SLE and underscored the therapeutic potential of B cell–targeted strategies, driving the development of next-generation monoclonal antibodies and T cell– redirecting approaches such as CAR T cells and bispecific T cell engagers^2^. While the long-term outcomes of these therapies remain under evaluation, emerging evidence suggests that resistance to B cell depletion may be rooted in the defect of innate immune mechanisms that sustain B cell activation. Therefore, although deeper or more sustained B-cell depletion may improve short-term outcomes, restoration of immune tolerance in SLE is likely to require concomitant targeting of innate inflammatory pathways.

SLE is associated with defects in multiple B cell subsets, including increased immature B cells, autoreactive plasma cells, and double-negative (IgD^−^CD27^−^) B cells, precursors of antibody-producing plasma cells^3^. This dysregulation is accompanied by a marked numerical and functional reduction in IL-10–producing regulatory B cells (Bregs), a population that suppresses inflammation and is associated with favourable outcomes^4^. This imbalance between effector and regulatory B cells is increasingly recognised as a hallmark of active disease in SLE. Mounting evidence indicates that B-cell abnormalities can arise downstream of aberrant neutrophil activation. Neutrophils are hyperactivated in SLE, with an expanded subset of low-density granulocytes prone to spontaneous release of neutrophil extracellular traps (NETs). Unlike canonical NETs, SLE-NETs are enriched in oxidised mitochondrial DNA (ox-mtDNA), which potently activates interferon-producing dendritic cells and the NLRP3 inflammasome^5,6^.

In lupus-prone mice, ox-mtDNA triggers anti-DNA antibody production and renal inflammation resembling lupus-nephritis^7^. We and others have shown that NETs are deposited in inflamed kidneys in immune-complex nephritis, and that inhibiting NET formation ameliorates disease in preclinical models^8,9^.Notably, patients treated with rituximab or CD19-targeted CAR-T cell therapies develop late-onset neutropenia that often resolves and coincides with B-cell repopulation suggesting bidirectional functional crosstalk between neutrophils and B cells^10,11^.

Mitochondria generate energy and maintain redox balance through oxidative phosphorylation (OXPHOS), which produces reactive oxygen species (ROS) as by-products. At homeostatic levels, mitochondrial ROS (mtROS) act as second messengers that regulate redox sensitive signalling pathways, but in excess they cause oxidative damage and metabolic dysfunction. ROS are detoxified mainly by the glutathione and thioredoxin systems, both of which depend on a constant supply of NADPH. Although the role of NADPH balance in B-cell differentiation is not yet defined, genetic studies suggest it is important as variants in NADPH-oxidase complex genes increase SLE risk^12^, and patients with chronic granulomatous disease, who lack functional NADPH-oxidase, often develop lupus-like symptoms^13^.

Our recent work shows that Breg differentiation depends on thioredoxin to keep ROS at physiological levels. When ROS levels exceed physiological thresholds, B-cells are pushed toward plasma-cell differentiation at the expense of Bregs^14^. Both thioredoxin recycling and cholesterol biosynthesis^15^ depend on NADPH availability. Under conditions of oxidative stress, increased antioxidant demand diverts NADPH away from cholesterol biosynthesis toward redox buffering. This metabolic shift disrupts lipid trafficking, causing mitochondrial cholesterol accumulation, reduced membrane fluidity, and impaired function^16^. Importantly, homeostatic cholesterol biosynthesis is essential for Breg IL-10 production via geranylgeranyl pyrophosphate^17^. Together, these findings suggest a metabolic trade-off in NADPH “allocation” between antioxidant defence and cholesterol synthesis that critically determines B cell differentiation.

Ox-mtDNA released in NETs places metabolic stress on B-cells. Following uptake, ox-mtDNA activates endosomal TLR9, cytosolic sensors such as cGAS^18^, and the inflammasome^19^. This triggers mitochondrial membrane instability and disrupts electron-transport chain activity^20^. Thus, we hypothesized that neutrophil-derived mtDNA drives B-cell metabolic dysfunction in SLE.

Here, we demonstrate how NETs and mitochondrial DNA released by SLE neutrophils modulate B cell responses by altering B cell mitochondrial and cholesterol metabolism and function promoting differentiation toward pro-inflammatory B cells while suppressing the development of IL-10^+^regulatory B cells.

### Single-cell profiling reveals proinflammatory neutrophil and B-cell subsets in lupus nephritis in peripheral blood and tissue

Neutrophils in SLE differ markedly from those in healthy individuals in both phenotype and function. A key feature of SLE is the expansion of low-density granulocytes (LDGs), a subset of neutrophils that co-purify with peripheral blood mononuclear cells (PBMCs) during density-gradient separation^21^. LDGs are expanded in the peripheral blood of patients with SLE, particularly in those with lupus nephritis, and have been shown to exhibit an immature phenotype and to undergo spontaneous NETosis^22–24^.

To further characterise LDG heterogeneity in SLE, we analysed the only publicly available single-cell RNA-sequencing dataset of LDGs derived from three patients with early renal involvement, no healthy controls were included in this dataset for comparison^22^. LDGs were identified based on expression of the neutrophil-specific genes ELANE and FCGR3B (Figure 1A and B). Dimensionality reduction using t-SNE revealed five distinct LDG clusters. Three clusters displayed transcriptional features of mature neutrophils, defined by expression of CD16b, whereas two clusters expressed high levels of ELANE, a primary granule gene characteristic of immature granulocytes (Figure C). Notably, immature LDGs selectively upregulated genes involved in NETosis, as well as pathways related to oxidative phosphorylation (OXPHOS), inflammasome activation, and interaction with the endothelium (Figure D and E).

**Figure 1.**
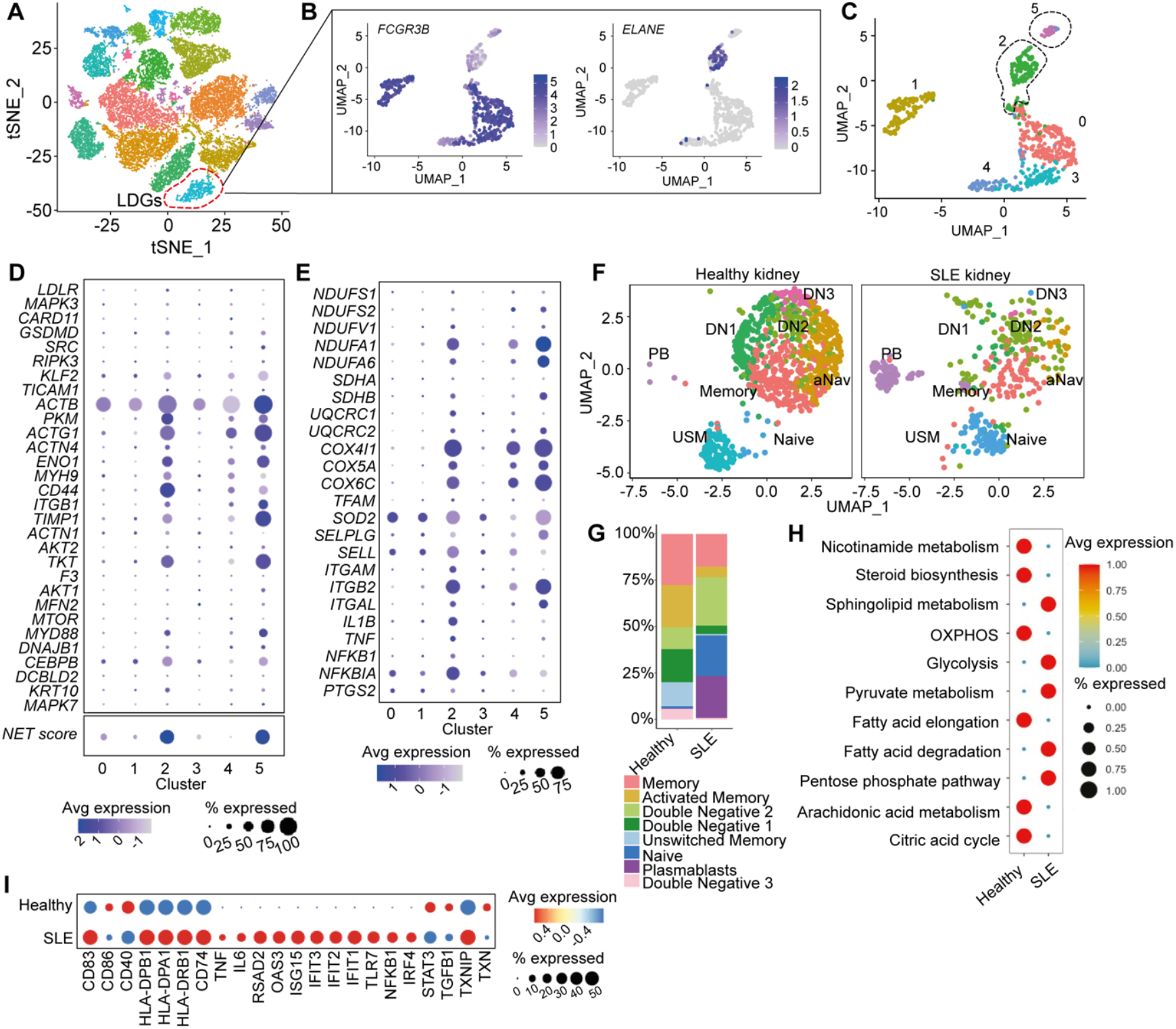
Single-cell profiling reveals proinflammatory neutrophil and B-cell subsets in lupus nephritis in peripheral blood and tissue. A. UMAP showing t-SNE plot representing gene expression in single cells from SLE-PBMCs (data from^22^). B. and C. Five transcriptionally LDG clusters identified based on their expression of FCGR3B/ELANE D. and E. Dot plot showing expression of genes involved in NETosis and the combined NET score^50^ derived from these genes across LDG clusters and of genes (E) involved in mitochondrial OXPHOS, endothelial interaction and proinflammatory signalling F. UMAP and G. stacked bar plots showing B-cell clusters from the integrated scRNA-seq datasets from lupus nephritis^26^ and healthy kidney^27^. H. Metabolic pathway analysis in kidney B cells from healthy and lupus nephritis kidneys. I. DotPlots of pro-inflammatory and regulatory gene expression and metabolic pathway analysis on healthy and lupus kidney B cells.

These data suggest that LDGs in SLE are a heterogeneous population, with an immature subset displaying heightened NETosis and the potential to activate the inflammasome and interact with inflamed endothelium.

Independent evidence for pathogenic neutrophil–B-cell crosstalk in SLE comes from GWAS studies, which identify genetic variants regulating neutrophil activation and innate immune signalling as key drivers of severe and treatment-refractory disease^25^.

While the contribution of B cells to SLE is well established, most transcriptomic and functional data highlighting abnormalities in B-cell composition, metabolism, and function are derived from peripheral blood. To determine whether findings in peripheral blood mirror those in kidney-resident B cells in SLE, the target organ where the most severe damage occurs, we leveraged single-cell transcriptomic datasets from patients with lupus nephritis (LN)^26^ and healthy controls^27^ to investigate the transcriptional and metabolic signatures of kidney-resident B cells in the context of SLE pathogenesis. Integration of these datasets revealed eight B-cell clusters (Supplementary Figure 1), including naïve B cells, class-switched and unswitched memory B cells, plasmablasts, and three subsets of double-negative (DN) B cells (DN1, DN2, and DN3). Compared with healthy kidneys, LN kidneys contained higher proportions of naïve B cells (CD27^lo^IGHD^hi^TCL1A^hi^FCER2^hi^), DN2 B cells (CD27^lo^IGHD^lo^CXCR5^lo^ZEB2^hi^TBX21^hi^), and plasmablasts (PRDM1^hi^XBP1^hi^) (Figure 1F-G). In SLE, naïve B cells are known to differentiate into pathogenic DN2 cells expressing T-bet and CD11c, which serve as precursors of autoreactive plasmablasts^3^. We therefore hypothesized that the expansion of DN2 B cells in the kidney may be accompanied by a reduction in regulatory B cells (Bregs).

Supporting this hypothesis, we observed that kidney-resident naïve B cells from LN patients expressed lower levels of Breg-associated genes, including TXN and TGFB1, compared with those from healthy controls. In addition, we found upregulation of TXNIP, an endogenous inhibitor of thioredoxin, in LN kidney B cells. Furthermore, LN B cells exhibited increased expression of pro-inflammatory markers involved in antigen presentation, Toll-like receptor engagement, nucleic acid and type I interferon sensing, as well as pro-inflammatory cytokines, including IL6 and TNF (Figure 1I).

We have previously shown that healthy Bregs depend on mitochondrial oxidative phosphorylation (OXPHOS) for their differentiation and suppressive function^14^. metabolic pathway analysis using the *scMetabolism* package^28^ revealed that LN kidney B cells exhibit reduced overall OXPHOS activity and increased glucose metabolism relative to healthy B cells (Figure 1H).

These findings suggest that SLE kidney B cells may be functionally deficient in OXPHOS, potentially impairing Breg differentiation and skewing them toward pro-inflammatory fates.

### Neutrophils from patients with active lupus nephritis induce mitochondrial oxidative stress and promote pro-inflammatory and plasmablast B-cell differentiation

We have shown that homeostatic levels of ROS are required for IL-10^+^Breg differentiation^14^. However, ROS in excess, induced by high concentrations of TLR ligands, suppressed IL-10^+^Bregs whilst promoting pro-inflammatory plasmablast differentiation. In SLE patients, B cells present with increased levels of mitochondrial ROS indicating defective OXPHOS which was accompanied by reduced IL-10^+^Breg frequencies. The source of the excessive ROS in SLE B cells is yet to be fully established. Our observation that high concentrations of TLR ligands suppressed Breg differentiation^14^ suggest that the high levels of nucleic acids, released primarily from neutrophils in SLE, activate TLRs in B cells and plasmacytoid dendritic cells (pDCs), thereby triggering NLRP3 inflammasome activity^5^ which may further amplify oxidative stress and inflammation.

To investigate this, we isolated polymorphonuclear neutrophils (PMNs) from patients with acute lupus nephritis (LN) and from healthy controls. Clinical and demographic patient characteristics are shown in Supplementary Table 1. Neutrophils were co-cultured with autologous or heterologous B cells for 72 hours. Healthy B cells co-cultured with SLE neutrophils exhibited increased mitochondrial ROS levels and mitochondrial membrane depolarisation compared with healthy B cells co-cultured with neutrophils from healthy donors (Figure 2A-B).

**Figure 2.**
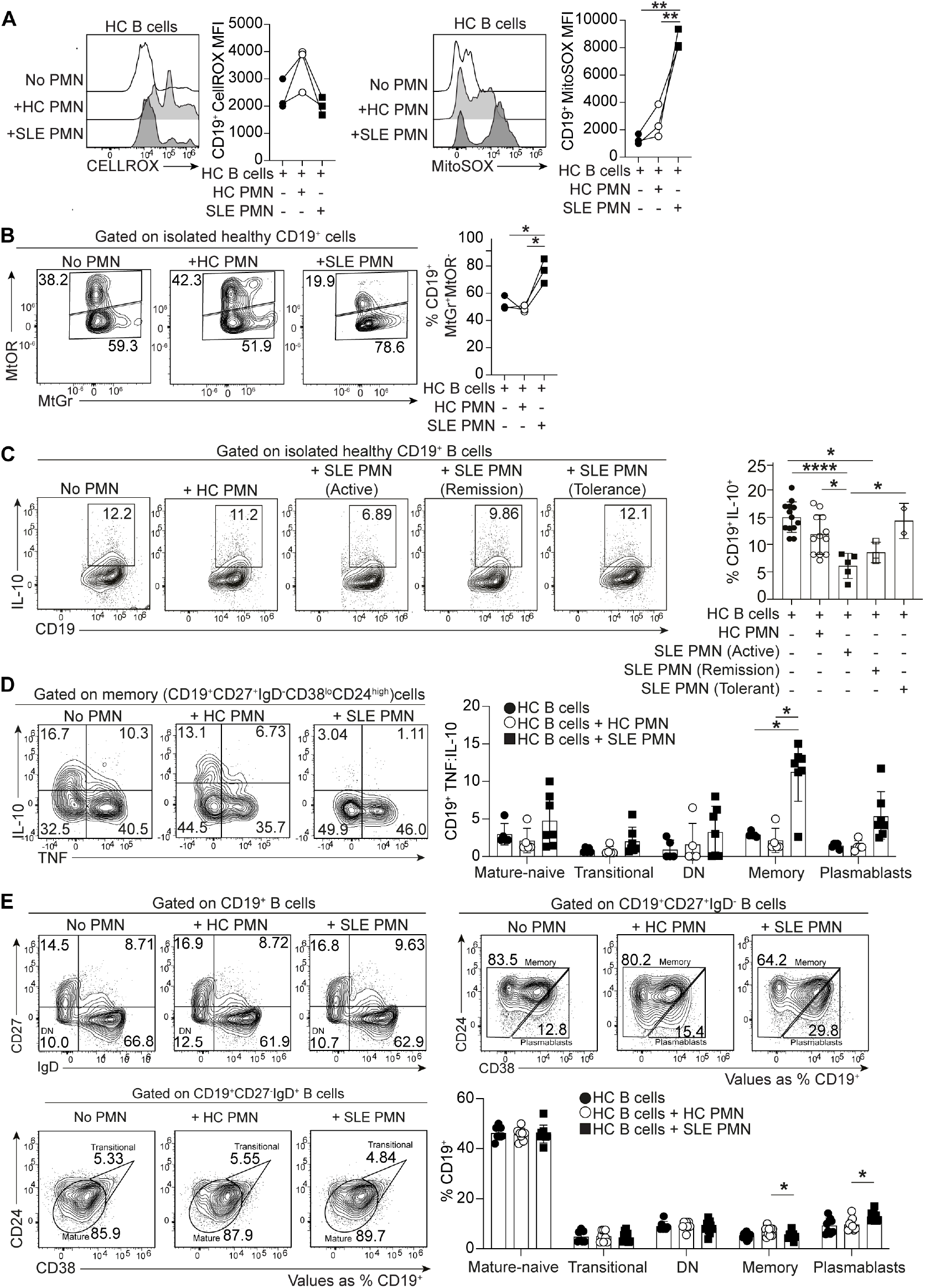
Neutrophils from patients with active lupus nephritis induce mitochondrial oxidative stress and promote pro-inflammatory and plasmablast B-cell differentiation Healthy or SLE CD19+ B-cells were stimulated with CpGC for 72h. A. Histograms and cumulative data show mitochondrial (MitoSOX) and cytosolic (CELL-ROX) ROS levels and B. mitochondrial membrane depolarisation (MitoGr^high^MitoOR^lo^) in healthy B cells co-cultured with neutrophils (PMNs) from healthy donors or SLE patients with lupus-nephritis, **p<0.01, by repeated-measures one-way ANOVA C. Representative contour plots and cumulative data show IL-10+CD19+ B-cell frequencies of healthy B cells co-cultured with PMNs from healthy donors or SLE patients with active disease, remission on maintenance immunosuppression, or disease tolerance.*p<0.05, ****p<0.0001 one-way ANOVA D. Representative contour plots and cumulative data show TNF:IL-10 expression ratios of CD19+ B cells from healthy donors co-cultured with healthy or SLE PMNs, repeated-measures one-way ANOVA, *p<0.05 Representative contour plots and cumulative data show frequencies of transitional (IgD+CD27-CD24hiCD38hi), mature naive (IgD+CD27-CD24hiCD38int), double-negative (DN, IgD+CD27-), memory (MBC, IgD-CD27+CD24+CD38lo) B cells and plasmablasts (PB, IgD-CD27+CD24-CD38^hi^), repeated-measures one-way ANOVA, *p<0.05

In parallel, neutrophils from patients with active lupus nephritis impaired IL-10^+^Breg differentiation in healthy B cells. In contrast, SLE neutrophils from patients in long-term tolerance, a rare patient cohort defined as individuals with normalised clinical and serological parameters who had discontinued immunosuppression without relapse for more than two years, promoted Breg differentiation. Approximately 5-10% of LN patients achieve disease tolerance^29^, whereas 45% of patients relapse after successful treatment of an initial episode. Notably, SLE neutrophils from patients in partial remission failed to fully restore Breg differentiation, suggesting that persistent neutrophil dysfunction, even in clinically improved patients, may predispose to disease relapse (Figure 2C).

In addition to impairing IL-10^+^Breg differentiation, SLE neutrophils skewed healthy B cells toward a pro-inflammatory phenotype, as evidenced by an increased TNF:IL-10 expression ratio in memory B cells (Figure 2D) and increased plasmablast frequencies (Figure 2E), compared with cultures containing healthy neutrophils.

Together, these findings demonstrate that neutrophils from patients with active lupus nephritis induce mitochondrial oxidative stress in B cells, impaired IL-10^+^Breg differentiation and a shift toward pro-inflammatory and plasmablast fates.

### Neutrophil extracellular trap–derived mitochondrial DNA suppresses IL-10^+^B-cell differentiation via TLR9 in lupus nephritis

We next determined whether the suppressive effect of SLE neutrophils on Breg differentiation was due to NET formation and TLR engagement. Healthy B cells were co-cultured with healthy or SLE neutrophils for 72h, with and without the TLR9 antagonist ODN 2088, which disrupts the colocalization of CpG ODNs with TLR9 in endosomal vesicles without affecting cellular binding and uptake. We also considered the ratio of TNF to IL-10 as a measure of B-effector versus B-regulatory function.

Compared to healthy B cells cultured alone, or with healthy neutrophils, healthy B cells cultured with SLE neutrophils presented with an increased TNF:IL-10 ratio. Healthy B cells cultured with SLE neutrophils in the presence of ODN 2088 showed a restored TNF:IL-10 ratio at similar levels to that of healthy B cells cultured with or without healthy neutrophils (Figure 3A). We then isolated SLE and healthy neutrophils, and incubated neutrophils overnight to induce spontaneous NETosis before harvesting supernatants containing NETs. Healthy B cells cultured with SLE neutrophil supernatants presented with reduced frequencies of IL-10^+^Bregs, compared to healthy B cells cultured with or without healthy neutrophils supernatants (Figure 3B). The addition of DNAse SLE neutrophil supernatants cultured with healthy B cells increased the frequencies of IL-10^+^Bregs, compared to those cultured in the absence of DNAse (Figure 3B). Together, these findings suggest that SLE neutrophils suppress Breg differentiation via NETs containing TLR ligands. We addressed whether SLE NETs suppress IL-10^+^Breg differentiation via mtDNA. We cultured healthy B cells with mtDNA or genomic DNA isolated from healthy or SLE neutrophils. Healthy B cells cultured with SLE neutrophil mtDNA showed reduced frequencies of IL-10+Bregs, compared to healthy B cells cultured with SLE neutrophil genomic DNA, healthy neutrophil mtDNA, or healthy neutrophil genomic DNA. There were no differences between any other culture conditions (Figure 3C).

**Figure 3.**
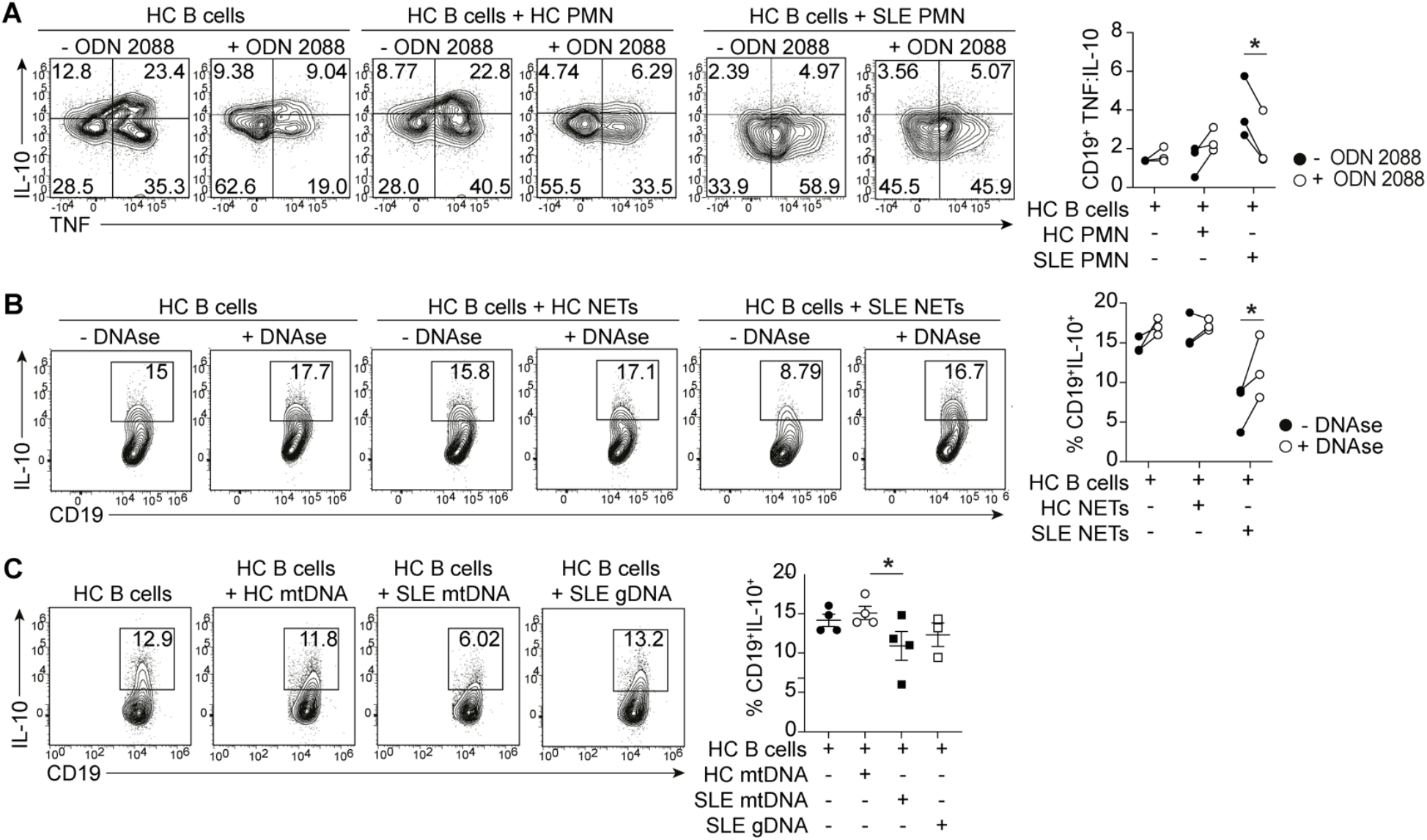
Neutrophil extracellular trap–derived mitochondrial DNA suppresses IL-10^+^ B-cell differentiation via TLR9 in lupus nephritis. A. Representative contour plots and cumulative data show TNF/IL-10 expression ratios of CD19+ B cells from healthy donors co-cultured with healthy or SLE PMNs, with or without the TLR9 agonist ODN 2088, \**P*<0.05, by repeated measures two-way ANOVA B. Representative contour plots and frequencies of IL-10+ CD19+ B-cells with supernatants from NETotic PMNs (healthy or SLE), with or without DNase I, \**P*<0.05,by repeated mesures two-way ANOVA, C. Representative contour plots and cumulative data show frequencies of CD19^+^IL-10^+^ B cells after 72h stimulation of healthy B cells with CpGC, co-cultured with or without mitochondrial or genomic DNA (gDNA) from healthy and SLE PMNs, \**P*<0.05, by repeated-measures one-way ANOVA

### Redox-driven NADPH imbalance is associated with altered cholesterol metabolism and regulatory B-cell differentiation in lupus nephritis

We have demonstrated that oxidative stress in SLE B cells is induced by NET-derived mitochondrial DNA, together with prior observations showing reduced thioredoxin expression in SLE B cells^14^. Nicotinamide adenine dinucleotide phosphate (NADPH) plays a central role in both antioxidant defence and de novo cholesterol biosynthesis. Under conditions of oxidative stress, NADPH is preferentially consumed to sustain antioxidant pathways, including thioredoxin recycling, limiting its availability for cholesterol synthesis^15,30^. We therefore hypothesise that oxidative stress in B cells, driven by NET-derived mitochondrial DNA, accelerates NADPH utilisation for redox buffering at the expense of *de novo* cholesterol biosynthesis.

We analysed integrated single-cell RNA-sequencing data from kidney B cells derived from healthy donors and patients with lupus nephritis and found that LN B cells downregulated genes involved in *de novo* cholesterol biosynthesis while upregulating genes associated with intracellular cholesterol trafficking and mitochondrial import (Figure 4A). While transcriptional signatures do not directly measure metabolic flux, they provide strong evidence for coordinated rewiring of redox and lipid pathways in LN B cells. We subsequently, validated these findings *in vitro*, using filipin staining, which binds unesterified free cholesterol, in *ex vivo* B cells isolated from the peripheral blood of patients with active lupus nephritis, patients in remission, and age-matched healthy controls. This analysis revealed a disease activity– dependent increase in intracellular unesterified cholesterol (Figure 4B). Notably, transitional and memory B cells, as well as plasmablasts from acute LN patients, exhibited significantly higher intracellular cholesterol levels compared with healthy controls.

**Figure 4.**
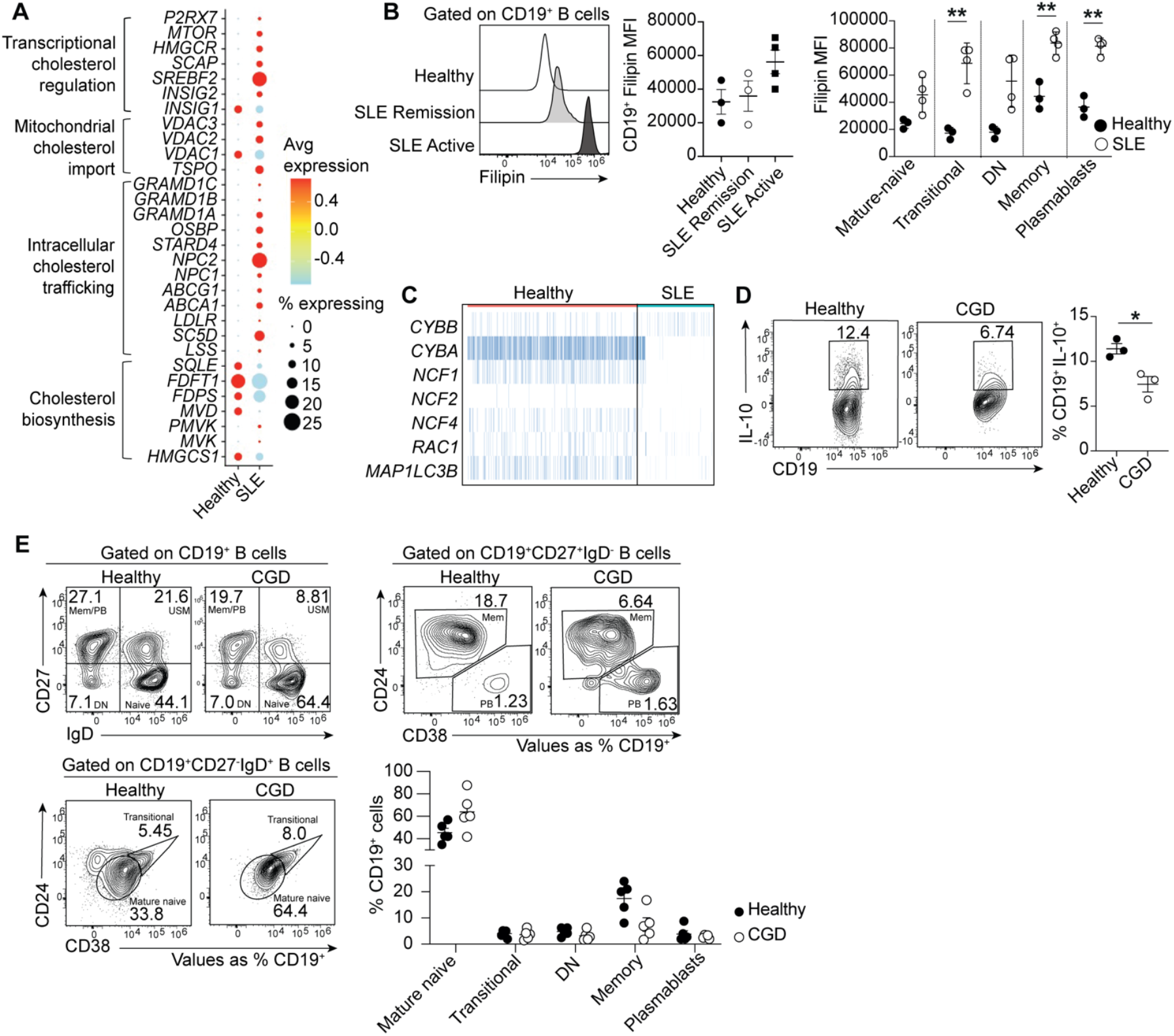
Redox-driven NADPH imbalance is associated with altered cholesterol metabolism and regulatory B-cell differentiation in lupus nephritis. A. Dot plot showing expression of genes involved in cholesterol biosynthesis and intracellular trafficking on the integrated scRNA seq datasets from lupus nephritis^26^ and healthy kidney^27^. B. Histogram and cumulative data showing Filipin MFI on ex vivo B cells from patients with SLE during active disease and in remission and healthy controls and Filipin MFI of B cell subsets in patients with active SLE and healthy controls, **P<0.01 by unpaired t-test. C. Heatmap showing expression of genes encoding NADPH complex subunits and LC3 in B cells from healthy and lupus nephritis kidneys from the integrated scRNA seq datasets from lupus nephritis^26^ and healthy kidney^27^. D. Representative contour plots and cumulative data show frequencies of CD19+IL-10+ B cells after 72h CpGC stimulation of B cells from healthy controls and CGD patients stimulated with CpGC for 72 hours, P<0.05 by unpaired t-test. E. Representative cumulative data show frequencies of transitional (IgD+CD27-CD24hiCD38hi), mature naive (IgD+CD27-CD24hiCD38int), double-negative (DN, IgD+CD27-), memory (MBC, IgD-CD27+CD24+CD38lo) B cells and plasmablasts (PB, IgD-CD27+CD24-CD38hi) in ex vivo peripheral blood of healthy controls and chronic granulomatous disease (CGD) patients., *P<0.05 by unpaired t-test.

NADPH is the sole substrate of the NADPH complex oxidase, a multi-subunit enzyme that, upon assembly, generates superoxide anion, which is subsequently converted into further reactive oxygen species (ROS) required for microbial killing and canonical NETosis. Polygenic variation in components of the NADPH oxidase complex have been linked to SLE, further highlighting the critical role of NADPH oxidase in disease susceptibility^31^.

In our integrated scRNA-seq data of immune cells from healthy and lupus nephritis kidneys, we observed that B cells from lupus nephritis patients downregulate genes encoding various subunits of the NADPH oxidase complex, as well as *MAP1LC3B*, which encodes LC3 (Figure 4C). As a proof of principle, to evaluate the functional consequences of NADPH deficiency on B cell differentiation, we analysed B cell subsets in patients with CGD carrying NADPH-associated genetic variants. Patients with CGD who are NADPH-deficient, exhibited an expanded naïve B cell compartment and reduced classical memory B cells consistent with previous reports^32,33^(Figure 4D). Furthermore, analysis of CD19^+^IL-10^+^B cells revealed a significant reduction in the differentiation of IL-10–producing regulatory B cells in NADPH-deficient patients compared to healthy controls (Figure 4E).

This data suggest that NET-derived mitochondrial DNA induces oxidative stress in SLE B cells, driving altered NADPH utilisation. This metabolic imbalance leads to altered lipid trafficking and the accumulation of cholesterol within plasma and mitochondrial membranes. NADPH deficiency is associated with impaired IL-10^+^regulatory B-cell differentiation, linking redox imbalance to B-cell dysregulation in lupus nephritis.

## Discussion

We show that neutrophils from patients with active lupus nephritis suppress IL-10–producing regulatory B-cell differentiation and promote pro-inflammatory and plasmablast differentiation through the release of NETs enriched in mitochondrial DNA. This effect is mediated by nucleic acid–dependent signalling, as both TLR9 inhibition and enzymatic degradation of NETs restore regulatory B-cell differentiation in vitro. These data support a model in which mitochondrial DNA contained within NETs represents a dominant upstream signal that perturbs B-cell metabolism and homeostasis in lupus, particularly within the renal microenvironment. Importantly, the ability of neutrophils from patients in long-term tolerance to restore Breg differentiation highlights neutrophil dysfunction as a dynamic and potentially reversible driver of immune dysregulation in SLE. Persistent neutrophil-mediated redox imbalance, even in partial clinical remission, may therefore represent a key mechanism predisposing to disease relapse and a potential therapeutic target.

In SLE, defective nucleic acid clearance leads to accumulation of endogenous TLR ligands, including NET-derived mtDNA, which potently activates endosomal TLRs. While TLR9 is traditionally associated with MyD88-dependent NFkB activation and inflammation, recent studies suggest that it may also signal via anti-inflammatory pathways in a context-dependent manner. Our data show that both TLR9 inhibition and DNAse treatment restore IL-10^+^Breg differentiation in the presence of SLE neutrophils suggesting that aberrant TLR9 signaling induced by NET-associated mtDNA disrupts Breg function. Although inhibitors of NET formation have shown efficacy in pre-clinical and clinical studies of neutrophil-driven diseases^34–36^, they have been largely ineffective in SLE. This may be due to their focus on canonical NETosis pathways, which rely on NADPH oxidase and genomic DNA. In contrast, NETs in SLE are enriched in mtDNA, and recent work has shown that targeting mtDNA release via inhibition of VDAC oligomerisation can improve outcomes in pre-clinical lupus models^37^.

These observations are supported by genome-wide association studies in SLE, which have identified numerous risk variants associated with disease development, particularly in genes involved in innate–adaptive immune system crosstalk. These include mutations in DNases responsible for degrading neutrophil extracellular traps, genes related to Toll-like receptor signalling, and those implicated in type I interferonopathies^38^.

The NADPH oxidase complex plays a critical role in shaping both neutrophil and B cell metabolism and function, thereby contributing to autoimmune pathogenesis. In the absence of functional NADPH activity, neutrophils release NETs enriched in oxidised mitochondrial DNA, which are more resistant to degradation and can trigger type I interferon (IFN-α) production by plasmacytoid dendritic cells^5,6^. Individuals with CGD, as well as female carriers, can develop autoimmune features, including discoid lupus and inflammatory arthritis^39–41^. Recent studies have also shown that NADPH activity regulates TLR ligand activation via LC3-mediated endosomal maturation and lack of NADPH complex deficient B cells exhibit enhanced signalling downstream of endosomal TLRs, increased humoral responses to nucleic acid-containing antigens, and the propensity toward humoral autoimmunity^42^. Our data support these findings by showing that SLE B cells in the lupus nephritis kidney downregulate genes encoding both NADPH oxidase subunits and LC3. Notably, the NADPH–cholesterol– mitochondrial axis has been implicated in IL-10^+^regulatory B cell differentiation^15,17^. Consistent with this, we observe reduced IL-10^+^Breg differentiation in CGD patients lacking functional NADPH oxidase compared to healthy controls.

This metabolic imbalance leads to altered lipid trafficking and the accumulation of cholesterol within plasma and mitochondrial membranes. Both thioredoxin recycling and cholesterol biosynthesis depend on NADPH, and diversion of NADPH toward antioxidant defence under oxidative stress reduces cholesterol synthesis, resulting in mitochondrial cholesterol accumulation, decreased membrane fluidity, and impaired mitochondrial function^16^. Moreover, cholesterol accumulation in mitochondrial and ER membranes alters their composition, impairing enzyme access and arachidonic acid release. This skews lipid mediators that are secreted by B-cells and can affect neutrophil activation towards pro-inflammatory leukotrienes, shown to be elevated in the urine of lupus patients^43^ at the expense of pro-resolving lipoxins^44^, supporting a bidirectional cross-talk between neutrophils-B cells in SLE.

To date, there is no cure for systemic lupus erythematosus (SLE), and current therapies primarily focus on B cell depletion. However, this strategy targets both effector B cells and regulatory B cells—a concern underscored during the COVID-19 pandemic, when immunosuppressed patients experienced increased infection-related mortality^45,46^. Pharmacological modulators of mitochondrial metabolism are currently being explored in neuroinflammatory diseases^47^ and cancer^48^, and have shown preclinical efficacy in models of lupus nephritis^37,49^. Based on our findings, we propose a paradigm shift: rather than eliminating B cells essential for immune homeostasis, therapeutic strategies should aim to correct dysregulated metabolic pathways to restore the balance between effector and regulatory B cells.

## Conclusion

Together, these findings support a model in which NET-derived mitochondrial DNA induces sustained oxidative stress in B cells, driving maladaptive NADPH allocation that disrupts cholesterol homeostasis, mitochondrial function, and regulatory B-cell differentiation. This neutrophil-driven metabolic reprogramming skews B-cell responses toward pathogenic effector states and may contribute to disease persistence and relapse, even in the setting of partial clinical remission. Targeting metabolic checkpoints that restore redox balance and lipid homeostasis, rather than indiscriminate B-cell depletion, may therefore represent a more precise strategy to re-establish immune regulation in lupus nephritis.

## Acknowledgements

This work was supported by the Rosetrees Trust, Lupus UK, and the 3TR (Taxonomy, Treatment, Targets and Remission) consortium. We thank the funders for their support and the patients and healthy donors who contributed samples to this study.

## Figure

**Supplementary Figure 1.**
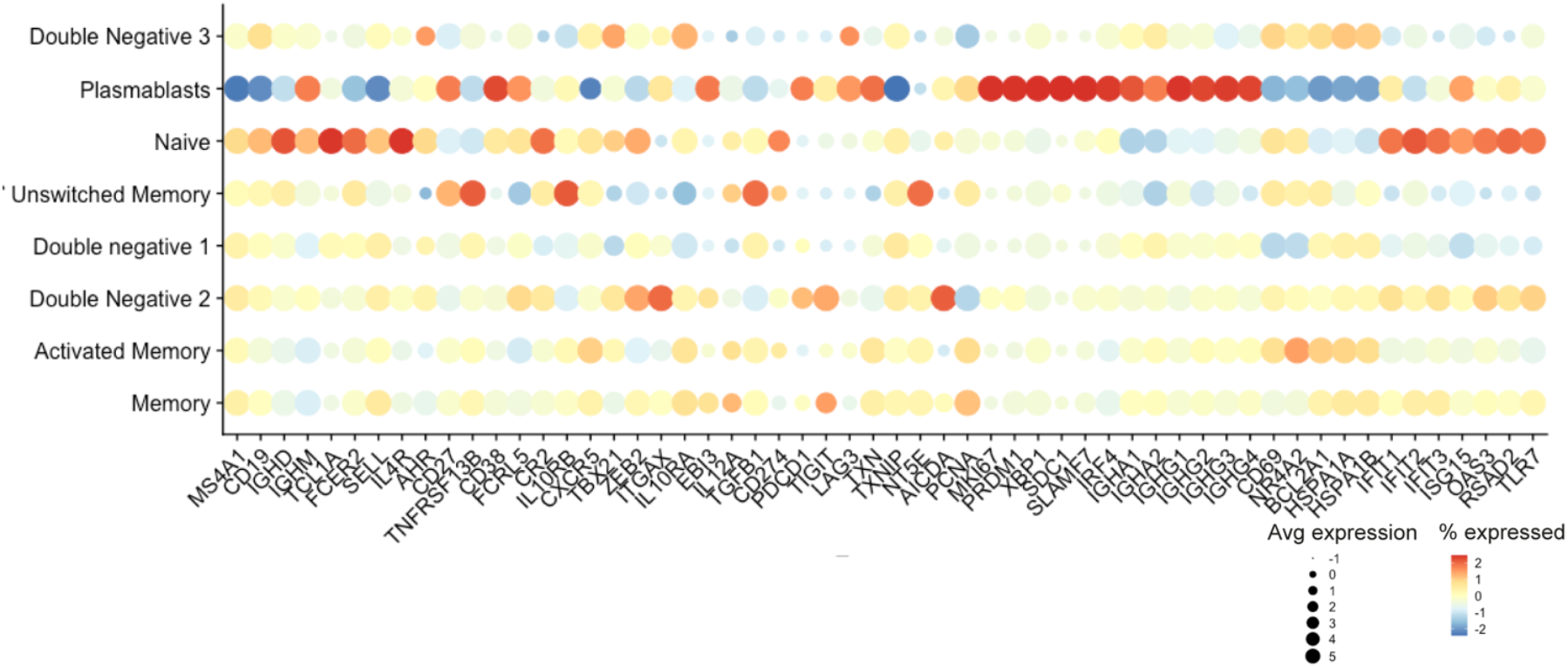
Dotplot showing for each B cell cluster as defined by selected key B cell markers in the integrated data sourced from lupus nephritis ^26^ and healthy kidney^27^ B cells.

**Supplementary Table 1.** Clinical and demographic characteristics of SLE patients.

Clinical and demographic characteristics of SLE patients
| Patient | Gender | Ethnicity | Age | uPCR<br>mg/mmol | eGFR<br>mL/min/<br>1.73m<sup>2</sup> | C3<br>g/L | C4<br>g/L | dsDNA | LN<br>class | Pred daily<br>dose/mg | MMF<br>daily<br>dose/mg |
| --- | --- | --- | --- | --- | --- | --- | --- | --- | --- | --- | --- |
| 1 | Female | Indian | 27 | 112 | >90 | 0.71 | 0.05 | 92 | IV | 30 | 2000 |
| 2 | Female | Black | 36 | 148 | >90 | 0.84 | 0.06 | 30 | IV+V | 5 | 1000 |
| 3 | Female | Black | 39 | 300 | 30 | 0.7 | 0.04 | 213 | III | 0 | 0 |
| 4 | Male | Black | 29 | 239 | 62 | 0.6 | 0.1 | 349 | IV | 15 | 2000 |
| 5 | Female | Black | 37 | 141 | >90 | 0.81 | 0.07 | 140 | IV | 5 | 1000 |
| 6 | Female | Black | 46 | 117 | 51 | 1.31 | 0.36 | 1.1 | III+V | 5 | 1000 |
| 7 | Female | White | 27 | 921 | 79 | 0.69 | 0.10 | 27 | IV | 40 | 1000 |
| 8 | Female | Black | 21 | 230 | 75 | 0.78 | 0.1 | 3.8 | V | 30 | 2000 |
| 9 | Male | Indian | 23 | 378 | 64 | 0.86 | 0.17 | 5.4 | III | 50 | 2000 |
| 10 | Female | Black | 25 | 268 | >90 | 0.79 | 0.22 | 268 | IV+V | 0 | 0 |
| 11 | Female | White | 44 | 717 | 82 | 1.79 | 0.33 | 1.3 | IV+V | 30 | 2000 |
| 12 | Female | Hispanic | 24 | 872 | >90 | 0.68 | 0.13 | 179 | IV+V | 10 | 2000 |
| 13 | Female | Asian | 36 | 128 | 45 | 0.91 | 0.19 | 0.7 | IV | 0 | 1000 |
| 14 | Male | Indian | 39 | 101 | 38 | 0.64 | 0.05 | 2.2 | IV | 0 | 500 |
| 15 | Male | Indian | 25 | 130 | 81 | 1.2 | 0.28 | 4.7 | IV | 0 | 1000 |
| 16 | Male | Indian | 45 | 7 | >90 | 1.09 | 0.17 | <0.6 | III | 0 | 500 |
| 17 | Female | White | 34 | 24 | >90 | 1.4 | 0.33 | 8.3 | IV | 0 | 500 |
| 18 | Female | Hispanic | 50 | 5 | >90 | 1.2 | 0.4 | 1 | III | 0 | 0 |
| 19 | Female | White | 46 | 10 | >90 | 1.5 | 0.3 | 3 | IV+V | 0 | 0 |

## Methods

### Patient selection

All included SLE patients had biopsy-proven class III or IV lupus nephritis. Patients were eligible if they had active disease (n=12) and were acute presentations either treatment-or steroid-naïve, minimally treated (≤1 g/day prednisolone), or had relapsing disease while receiving low-dose immunosuppression (prednisolone ≤5 mg/day, azathioprine ≤2 mg/kg/day, or mycophenolate mofetil ≤1 g/day). Clinical remission (n=5) was defined as a >50% reduction in proteinuria with stable or improved estimated glomerular filtration rate (eGFR) at 6 months. Immunological tolerance (n=2) was defined as sustained remission off all immunosuppressive therapy for >2 years without relapse. Patients who had received cyclophosphamide within the preceding 12 months were excluded.

### Sample collection and B cell/neutrophil isolation

40ml Peripheral blood was collected from LN patients attending the Royal Free hospital outpatient lupus clinic (n=19) and age/sex-matched healthy donors(n=13). Blood was diluted 1:1 in RPMI 1640, and PBMC monolayers were isolated using STEMCELL SepMate density gradient based centrifugation. B cells were isolated using the STEMCELL EasySep B cell enrichment kit (STEMCELL, 19054) according to manufacturers instructions. Neutrophils were isolated using EasySep™ Direct Human Neutrophil Isolation Kit Catalog # 19666.

### Cell culture assays

Isolated B cells and neutrophils were seeded at a density of 1 × 10^6^ cells/mL in round-bottom 96-well plates and cultured for 72 h with CpG-C ODN 2395 (1 μM; InvivoGen, cat. no. tlrl-2395-1) in RPMI-1640 containing L-glutamine (Sigma-Aldrich). Culture media were supplemented with 10% fetal calf serum (FCS) (LabTech) and 1% penicillin/streptomycin (100 U/mL penicillin and 100 μg/mL streptomycin; Sigma-Aldrich). Where indicated, cultures were co-treated with the TLR9 antagonist ODN 2088 (InvivoGen, cat. no. tlrl-2088) for 72 h at 37 °C in 5% CO_2_.

Total cellular and mitochondrial reactive oxygen species (ROS) were quantified using CellROX and MitoSOX assays, respectively (Thermo Fisher Scientific; cat. nos. C10443 and M36008), according to the manufacturer’s instructions.

### NET and mitochondrial DNA generation

For NET isolation, neutrophils were incubated overnight at 37 °C in 5% CO_2_ to allow spontaneous NET formation. Cultures were subsequently treated with or without DNase I (10 U/mL). Neutrophil supernatants were then harvested and added to B-cell cultures.

Mitochondrial DNA (mtDNA) was isolated using the Mitochondrial DNA Isolation Kit (Abcam, cat. no. ab65321) according to the manufacturer’s instructions.

### Flow cytometry

Single cell suspensions were stained with LIVE/DEAD™ Fixable Blue Dead Cell Stain (ThermoFisher) at 1:500 in PBS for 20 minutes at room temperature. Surface markers were stained in PBS (2% FCS) for 20 minutes at 4°C. For intracellular staining, cells were fixed for 20 minutes in paraformaldehyde (ThermoFisher Scientific, 88-8824-00), followed by permeabilisation for 5 minutes (ThermoFisher, 88-8824-00). Intracellular antibodies were incubated for 40 minutes at 4°C. Data were acquired on an Aurora Spectral flow cytometer (Cytek) using spectral unmixing and subtraction of autofluorescence based on matched unstained controls. Analysis was performed using FlowJo (TreeStar).

### scRNA-sequencing analysis

Healthy and LN kidney biopsy scRNA-seq datasets were obtained from Arazi et al., and Stewart et al^1,2^. The raw transcript count matrices were loaded into R using RStudio and the Seurat package. Datasets were merged into a single Seurat object using the Harmony package. Output matrices were filtered to exclude cells with low gene frequencies (<300), high frequencies (>6000) or high percentages of mitochondrial DNA (>10%). Feature expression measurements for each cell were normalised by the total expression, multiplied by a scaling factor of 10,000, and log-normalised. Cell cycle regression was performed on the normalised dataset, and regressed data were scaled so each gene had a mean expression of 0 and variance of 1 across cells. Uniform Manifold Approximation and Projection (UMAP) was performed to cluster cells. B cells were subsetted based on expression of phenotypic markers *MS4A1, CD19, CD79A, IGJ, IGHG1*, and *SDC1* and LDG neutrophils based on expression of ELANE and FCGR3B

## Notes

### Competing Interest Statement

The authors have declared no competing interest.

## References

1. Almaani S, Meara A, Rovin BH: Update on Lupus Nephritis. CJASN 12: 825–835, 2017

2. Stockfelt M, Teng YKO, Vital EM: Opportunities and limitations of B cell depletion approaches in SLE. Nat Rev Rheumatol 21: 111–126, 2025

3. Jenks SA, Cashman KS, Zumaquero E, Marigorta UM, Patel AV, Wang X, et al.: Distinct Effector B Cells Induced by Unregulated Toll-like Receptor 7 Contribute to Pathogenic Responses in Systemic Lupus Erythematosus. Immunity 49: 725-739.e6, 2018

4. Blair PA, Noreña LY, Flores-Borja F, Rawlings DJ, Isenberg DA, Ehrenstein MR, et al.: CD19(+)CD24(hi)CD38(hi) B cells exhibit regulatory capacity in healthy individuals but are functionally impaired in systemic Lupus Erythematosus patients. Immunity 32: 129–140, 2010

5. Lood C, Blanco LP, Purmalek MM, Carmona-Rivera C, De Ravin SS, Smith CK, et al.: Neutrophil extracellular traps enriched in oxidized mitochondrial DNA are interferogenic and contribute to lupus-like disease. Nat. Med. 22: 146–153, 2016

6. Wu T, Chen X, Fan J, Ye P, Zhang J, Wang Z, et al.: Oxidative stress-induced release of mitochondrial DNA (mtDNA) promotes the progression of vitiligo by activating the cGAS-STING signaling pathway in monocytes. Free Radical Biology and Medicine 235: 43–55, 2025

7. Xian H, Watari K, Ohira M, Brito JS, He P, Onyuru J, et al.: Mitochondrial DNA oxidation propagates autoimmunity by enabling plasmacytoid dendritic cells to induce TFH differentiation. Nat Immunol 26: 1168–1181, 2025

8. Antonelou M, Michaëlsson E, Evans RDR, Wang CJ, Henderson SR, Walker LSK, et al.: Therapeutic Myeloperoxidase Inhibition Attenuates Neutrophil Activation, ANCA-Mediated Endothelial Damage, and Crescentic GN. JASN [Internet] 2019 Available from: https://jasn.asnjournals.org/content/early/2019/12/25/ASN.2019060618 [cited 2020 Jan 31]

9. Antonelou M, Evans RDR, Henderson SR, Salama AD: Neutrophils are key mediators in crescentic glomerulonephritis and targets for new therapeutic approaches. Nephrol Dial Transplant 37: 230–238, 2022

10. Wolach O, Bairey O, Lahav M: Late-onset neutropenia after rituximab treatment: case series and comprehensive review of the literature. Medicine (Baltimore) 89: 308–318, 2010

11. Neelapu SS, Tummala S, Kebriaei P, Wierda W, Locke FL, Lin Y, et al.: Toxicity management after chimeric antigen receptor T cell therapy: one size does not fit “ALL.” Nat Rev Clin Oncol 15: 218–218, 2018

12. Yokoyama N, Kawasaki A, Matsushita T, Furukawa H, Kondo Y, Hirano F, et al.: Association of NCF1 polymorphism with systemic lupus erythematosus and systemic sclerosis but not with ANCA-associated vasculitis in a Japanese population. Sci Rep 9: 16366, 2019

13. Cale CM, Morton L, Goldblatt D: Cutaneous and other lupus-like symptoms in carriers of X-linked chronic granulomatous disease: incidence and autoimmune serology. Clin Exp Immunol 148: 79–84, 2007

14. Bradford HF, McDonnell TCR, Stewart A, Skelton A, Ng J, Baig Z, et al.: Thioredoxin is a metabolic rheostat controlling regulatory B cells. Nat Immunol 25: 873–885, 2024

15. Bonora M, Morganti C, van Gastel N, Ito K, Calura E, Zanolla I, et al.: A mitochondrial NADPH-cholesterol axis regulates extracellular vesicle biogenesis to support hematopoietic stem cell fate. Cell Stem Cell 31: 359-377.e10, 2024

16. Solsona-Vilarrasa E, Fucho R, Torres S, Nuñez S, Nuño-Lámbarri N, Enrich C, et al.: Cholesterol enrichment in liver mitochondria impairs oxidative phosphorylation and disrupts the assembly of respiratory supercomplexes. Redox Biol 24: 101214, 2019

17. Bibby JA, Purvis HA, Hayday T, Chandra A, Okkenhaug K, Rosenzweig S, et al.: Cholesterol metabolism drives regulatory B cell IL-10 through provision of geranylgeranyl pyrophosphate. Nat Commun 11: 3412, 2020

18. Ghincea A, Woo S, Yu S, Pivarnik T, Fiorini V, Herzog EL, et al.: Mitochondrial DNA-Sensing Pathogen Recognition Receptors in Systemic Sclerosis-Associated Interstitial Lung Disease: a Review. Curr Treat Options in Rheum 9: 204–220, 2023

19. Xian H, Watari K, Sanchez-Lopez E, Offenberger J, Onyuru J, Sampath H, et al.: Oxidized DNA fragments exit mitochondria via mPTP- and VDAC-dependent channels to activate NLRP3 inflammasome and interferon signaling. Immunity 55: 1370-1385.e8, 2022

20. Giordano L, Ware SA, Lagranha CJ, Kaufman BA: Mitochondrial DNA signals driving immune responses: Why, How, Where? Cell Commun Signal 23: 192, 2025

21. Carmona-Rivera C, Kaplan MJ: Low density granulocytes: a distinct class of neutrophils in systemic autoimmunity. Semin Immunopathol 35: 455–463, 2013

22. Mistry P, Nakabo S, O’Neil L, Goel RR, Jiang K, Carmona-Rivera C, et al.: Transcriptomic, epigenetic, and functional analyses implicate neutrophil diversity in the pathogenesis of systemic lupus erythematosus. Proc Natl Acad Sci U S A 116: 25222–25228, 2019

23. Rahman S, Sagar D, Hanna RN, Lightfoot YL, Mistry P, Smith CK, et al.: Low-density granulocytes activate T cells and demonstrate a non-suppressive role in systemic lupus erythematosus. Annals of the Rheumatic Diseases 78: 957–966, 2019

24. Midgley A, McLaren Z, Moots RJ, Edwards SW, Beresford MW: The role of neutrophil apoptosis in juvenile-onset systemic lupus erythematosus. Arthritis Rheum 60: 2390–2401, 2009

25. Laurynenka V, Harley JB: The 330 risk loci known for systemic lupus erythematosus (SLE): a review. Front. Lupus [Internet] 2: 2024 Available from: https://www.frontiersin.org/journals/lupus/articles/10.3389/flupu.2024.1398035/full [cited 2026 Feb 3]

26. Arazi A, Rao DA, Berthier CC, Davidson A, Liu Y, Hoover PJ, et al.: The immune cell landscape in kidneys of patients with lupus nephritis. Nat Immunol 20: 902–914, 2019

27. Stewart BJ, Ferdinand JR, Young MD, Mitchell TJ, Loudon KW, Riding AM, et al.: Spatiotemporal immune zonation of the human kidney. Science 365: 1461–1466, 2019

28. Wu Y, Yang S, Ma J, Chen Z, Song G, Rao D, et al.: Spatiotemporal Immune Landscape of Colorectal Cancer Liver Metastasis at Single-Cell Level. Cancer Discov 12: 134–153, 2022

29. Alenzi F, Ateka-Barrutia O, Ken Cheah C, Khamashta M, Sangle SR, D’Cruz DP: Lupus Nephritis Outcomes after Stopping Immunosuppression. J Clin Med 13: 2211, 2024

30. Sies H, Jones DP: Reactive oxygen species (ROS) as pleiotropic physiological signalling agents. Nat Rev Mol Cell Biol 21: 363–383, 2020

31. Thomas DC: How the phagocyte NADPH oxidase regulates innate immunity. Free Radical Biology and Medicine 125: 44–52, 2018

32. Bleesing JJ, Souto-Carneiro MM, Savage WJ, Brown MR, Martinez C, Yavuz S, et al.: Patients with Chronic Granulomatous Disease Have a Reduced Peripheral Blood Memory B Cell Compartment1. The Journal of Immunology 176: 7096–7103, 2006

33. Pozo-Beltrán CF, Suárez-Gutiérrez MA, Yamazaki-Nakashimada MA, Medina-Vera I, Saracho-Weber F, Macías-Robles AP, et al.: B subset cells in patients with chronic granulomatous disease in a Mexican population. Allergologia et Immunopathologia 47: 372–377, 2019

34. Wigerblad G, Kaplan MJ: Neutrophil extracellular traps in systemic autoimmune and autoinflammatory diseases. Nat Rev Immunol 23: 274–288, 2023

35. Volpi C, Fallarino F, Pallotta MT, Bianchi R, Vacca C, Belladonna ML, et al.: High doses of CpG oligodeoxynucleotides stimulate a tolerogenic TLR9–TRIF pathway. Nat Commun 4: 1852, 2013

36. Leibler C, John S, Elsner RA, Thomas KB, Smita S, Joachim S, et al.: Genetic dissection of TLR9 reveals complex regulatory and cryptic proinflammatory roles in mouse lupus. Nat Immunol 23: 1457–1469, 2022

37. Kim J, Gupta R, Blanco LP, Yang S, Shteinfer-Kuzmine A, Wang K, et al.: VDAC oligomers form mitochondrial pores to release mtDNA fragments and promote lupus-like disease. Science 366: 1531–1536, 2019

38. Vinuesa CG, Shen N, Ware T: Genetics of SLE: mechanistic insights from monogenic disease and disease-associated variants. Nat Rev Nephrol 19: 558–572, 2023

39. Marciano BE, Zerbe CS, Falcone EL, Ding L, DeRavin SS, Daub J, et al.: X-linked carriers of chronic granulomatous disease: Illness, lyonization, and stability. J Allergy Clin Immunol 141: 365–371, 2018

40. Battersby AC, Braggins H, Pearce MS, Cale CM, Burns SO, Hackett S, et al.: Inflammatory and autoimmune manifestations in X-linked carriers of chronic granulomatous disease in the United Kingdom. Journal of Allergy and Clinical Immunology 140: 628-630.e6, 2017

41. Chiriaco M, Salfa I, Ursu GM, Cifaldi C, Di Cesare S, Rossi P, et al.: Immunological Aspects of X-Linked Chronic Granulomatous Disease Female Carriers. Antioxidants (Basel) 10: 891, 2021

42. Liu S, Lagos J, Shumlak NM, Largent AD, Lewis STE, Holder U, et al.: NADPH oxidase exerts a B cell–intrinsic contribution to lupus risk by modulating endosomal TLR signals. Journal of Experimental Medicine 221: e20230774, 2024

43. Hackshaw KV, Voelkel NF, Thomas RB, Westcott JY: Urine leukotriene E4 levels are elevated in patients with active systemic lupus erythematosus. J Rheumatol 19: 252–258, 1992

44. Podstawka J, Sinha S, Hiroki CH, Sarden N, Granton E, Labit E, et al.: Marginating transitional B cells modulate neutrophils in the lung during inflammation and pneumonia. J Exp Med 218: e20210409, 2021

45. Rutherford MA, Scott J, Karabayas M, Antonelou M, Gopaluni S, Gray D, et al.: Risk Factors for Severe Outcomes in Patients With Systemic Vasculitis and COVID-19: A Binational, Registry-Based Cohort Study. Arthritis Rheumatol 73: 1713–1719, 2021

46. Akhlaq A, Aamer S, Hasan KM, Muzammil TS, Sohail AH, Quazi MA, et al.: Systemic lupus erythematosus is associated with increased risk of mortality and acute kidney injury in patients with COVID-19 hospitalization: Insights from a National Inpatient Sample analysis. Lupus 33: 248–254, 2024

47. Xu J, Du W, Zhao Y, Lim K, Lu L, Zhang C, et al.: Mitochondria targeting drugs for neurodegenerative diseases—Design, mechanism and application. Acta Pharm Sin B 12: 2778–2789, 2022

48. Sainero-Alcolado L, Liaño-Pons J, Ruiz-Pérez MV, Arsenian-Henriksson M: Targeting mitochondrial metabolism for precision medicine in cancer. Cell Death Differ 29: 1304–1317, 2022

49. Blanco LP, Pedersen HL, Wang X, Lightfoot YL, Seto N, Carmona-Rivera C, et al.: Improved Mitochondrial Metabolism and Reduced Inflammation Following Attenuation of Murine Lupus With Coenzyme Q10 Analog Idebenone. Arthritis Rheumatol 72: 454–464, 2020

50. McDonald PC, Topham JT, Awrey S, Tavakoli H, Carroll R, Brown WS, et al.: Neutrophil extracellular trap gene expression signatures identify prognostic and targetable signaling axes for inhibiting pancreatic tumour metastasis. Commun Biol 8: 1006, 2025

